# EEG-based classification models reveal differential neural processing of words and images

**DOI:** 10.64898/2026.03.16.712233

**Authors:** Neda R. Morakabati, Alison S. Thiha, Eitan Schechtman

**Author notes:** **Corresponding author:** Eitan Schechtman.

## Abstract

**Background:** Machine learning methods employing neuroimaging data are useful for monitoring the activation of neural representations. Specifically, they can be used to discern the brain networks engaged in processing specific categories of items. This approach has been employed on neuroimaging data, including functional magnetic resonance imaging data and electroencephalography (EEG) data.

**New method:** Here, we present a task and an analytical pipeline for investigating category representations using EEG. Participants (*N* = 30) viewed a series of images and words of objects belonging to five categories (Animals, Tools, Food, Scenes, and Vehicles) and responded when items from the same category were presented consecutively.

**Results:** We trained support vector machines on EEG data within participants and found that both image trials and word trials yielded significant category classification accuracy, with image trials achieving higher accuracy than word trials. When comparing categories in a pair-wise fashion, all pairs were statistically distinguishable for image trials, whereas only one pair was distinguishable for word trials. Parietal and Left Temporal electrodes contributed more to image classification than Frontal and Right Temporal electrodes. Category-specific activity patterns also generalized across participants for image trials.

**Comparison with existing methods:** Our data and analytic pipeline yielded high classification accuracies, primarily for image trials, providing support for the utility of EEG data for neural decoding.

**Conclusions:** These methods can be instrumental for exploring the activation and reactivation of neural representations at the category level during wakefulness and, potentially, during offline states.

## 1 INTRODUCTION

Categorization is fundamental to how the mind and brain organize semantic knowledge about the world. Children learn to categorize objects and actions as early as infancy (Owen & Barnes, 2021), and it remains an efficient way for processing information throughout life. Over years of learning, category assignment comes easily and mitigates the cognitive load of new learning. Distinct categories are represented by separable neural networks, with more similar categories (e.g., dogs and cows) having closer representation in the brain relative to more distinct items (e.g., dogs and umbrellas), which have more distinguishable representations (Bracci & Op de Beeck, 2016; Kriegeskorte et al., 2008). Research on categorical representations in the brain has revealed multiple category-specific regions, including areas that are dedicated to the processing of faces, houses, scenes, and tools (Epstein & Kanwisher, 1998; Ishai et al., 1999; Lewis, 2006).

Historically, research on category representation in the brain relied primarily on lesion studies (see Gainotti, 2000, for review). Since the 1990s, studies have employed neuroimaging techniques that capture activity throughout the brain during category processing, with most early studies using functional magnetic resonance imaging (fMRI; e.g., Ishai et al., 1999). In this study, we examined the fidelity of categorical representation using scalp electroencephalography (EEG), leveraging its high temporal resolution. Classically, studies employing EEG to examine category selectivity have used event-related potentials (ERP), the time-locked neural response in a single electrode, averaged over multiple trials. For example, Bentin and colleagues discovered the N170 ERP, a negative deviation peaking ∼170 ms after the visual presentation of a face (Bentin et al., 1996). ERP analyses commonly specify a topographical location on the scalp and a specific latency at which effects are most pronounced. However, this approach does not facilitate the detection of nuanced (yet consistent) differences that are distributed across the scalp, potentially changing dynamically over time. For example, a slight lateral shift in topography may be lost when examining any single electrode due to signal-to-noise issues, but could be reliably identified when considering the pattern in a holistic way.

In recent years, methods were developed to harness these scalp-wide dynamics for examining neural representations across time and space. To achieve this, data are typically submitted to machine learning algorithms that are trained to distinguish between classes of stimuli, actions, and internal states. For example, Bae & Luck (2018) trained classification models on EEG data across electrodes to reveal covert shifts in spatial attention. Essentially, the models examined data from all electrodes at different time points and across different attentional states to extract information regarding spatial attention. Classification accuracy was taken as a proxy for the information reflected in the neural signal, suggesting the enactment of selective neural representation. Other studies have used similar approaches to more directly examine the distinct neural representations for perceived categories (e.g., Schreiner et al., 2021). Notably, the methods used in these studies were agnostic about which electrodes contributed most, instead leveraging a data-driven approach to identify neural patterns with minimal experimenter intervention.

In our work, we build on these recent methodological advances and use support vector machines (SVMs) on high-density scalp EEG data to demonstrate EEG-decodable representation across different categories and modalities. Participants came into the lab and viewed a series of alternating images and words representing objects from five categories: Animals, Tools, Food, Scenes, and Vehicles. Participants were instructed to indicate when the currently presented item was of the same category as the one immediately preceding it, while EEG data was collected. We trained SVMs on this data to classify the presented category and modality. All SVMs were trained and tested within-subjects, except for one analysis, which trained them across subjects to identify generalizable category-specific neural patterns. All code, stimuli, and data are publicly shared to enable further exploration of category representation in the brain.

## 2 METHODS

### 2.1 Participants

A total of 30 participants (19 women, 11 men) were recruited for this study, with a mean age of 22.5 ± 2.86 SD years (range 18-31). Of these participants, 18 were native English speakers, and the rest were non-native fluent speakers. The majority of participants were right-handed (*n* = 26), three were left-handed, and one was ambidextrous. The majority of our sample identified as Asian (*n* = 20), followed by White (*n* = 8) and Black/African American (*n* = 1), with one participant selecting “Not Listed / Decline to State.” Two people identified as Hispanic/Latino. Participants completed the study for either course credits or monetary compensation, and were recruited through flyers, emails, and social media platforms. Eligibility criteria required participants to be between 18 and 35 years of age, be fluent in English, and have no history of neurological disorders. Participants were instructed to abstain from using recreational drugs and alcohol for 24 hours prior to their lab appointment. On the day of the experiment, participants completed a consent form and a mental health questionnaire before being fitted with an EEG cap. The study protocol was reviewed and approved by the Institutional Review Board at the University of California, Irvine.

### 2.2 Materials

A total of 100 stimuli were used in this experiment, comprising 50 images and 50 matched words. Stimuli belonged to five categories: Animals, Tools, Food, Scenes, and Vehicles (Figure 1a). The same ten items were represented in word-and image-form for each category (e.g., for the animals category, the word “DOG” and an image of a dog were used, respectively). The image dimensions were 800 x 800 pixels, and they were sourced from the THINGS database (Hebart et al., 2019), the BOLD5000 database (Chang et al., 2019), and from online websites. The number of letters per word did not significantly differ across categories (*F*(4,45) = 1.96, *p* = 0.12; Animals: 4.2 ± 1.72 SD, Tools: 6.3 ± 2.53, Food: 6.8 ± 1.89, Scenes: 6.1 ± 1.97, Vehicles: 6.4 ± 2.61). Similarly, words did not differ in the number of graphemes (*F*(4,45) = 1.91, *p* = 0.13) or their Kučera-Francis written frequency (*F*(4,38) = 0.93, *p* = 0.46; calculated based on 43/50 words for which norms were available using the well-established resource; Kučera & Francis, 1967). All words were initially presented along with a corresponding image to avoid confusion or idiosyncratic interpretations and to maximize levels of concreteness and imaginability. Nevertheless, some differences in concreteness and imaginability may have still existed among the presented words, and the results should be interpreted with these limitations in mind.

**Figure 1:**
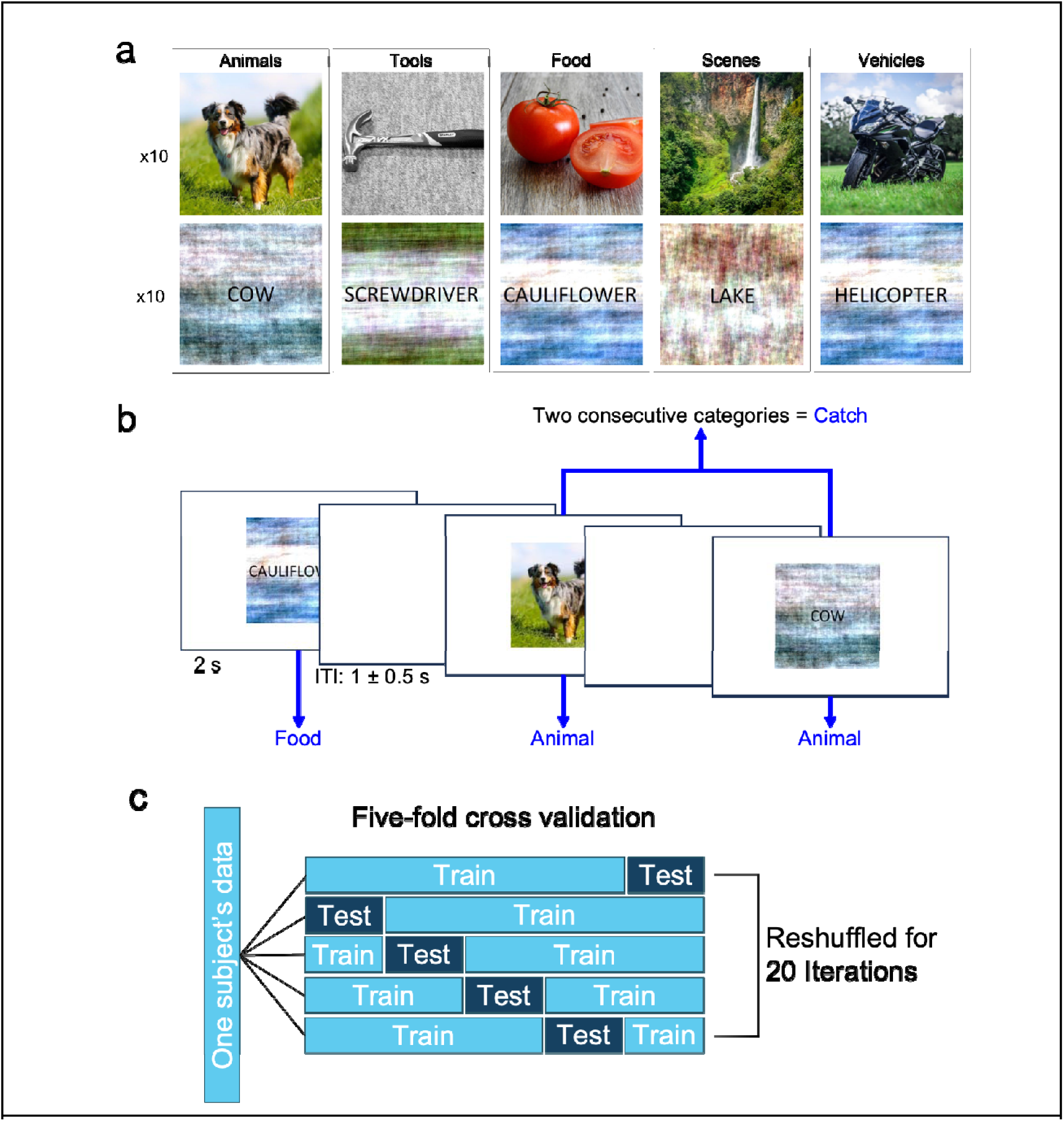
An overview of the materials, behavioral task, and analyses. **a**. Each category included ten images and ten words, totaling 100 items. Word stimuli were presented on a background matching the visual dimensions of the images. **b.** To elicit categorical brain activity, participants viewed an alternating series of images and words. Each item was presented for 2 s, followed by a blank screen for 0.5-1.5 s. Participants were instructed to press the spacebar when two items of the same category appeared in sequence. **c.** We used machine learning models to identify distinguishable activity (e.g., between categories) in our EEG data. Our analysis used support vector machines. Models employed five-fold cross-validation (i.e., each model was trained on 80% of the data and tested on the remaining 20%, repeated five times). Data was reshuffled over 20 iterations.

Both stimulus types (images and words) were included to ensure that the elicited neural responses corresponded to the conceptual meaning of each stimulus rather than merely distinguishing between images and words based on their physical appearance. Additionally, to control for potential confounds related to visual characteristics, each word stimulus was presented on an 800 x 800 pixel abstract background, which was a scrambled version of an image from the BOLD5000 database. The backgrounds were intended to match the physical features of the presented images.

### 2.3 EEG Setup and Application

Participants were fitted with a 64-channel EEG cap (ANT Neuro, Netherlands) with electrodes positioned in accordance with the standardized 10-20 system for EEG measurements. The two mastoid electrodes served as reference electrodes, and EOG activity was measured using two electrodes positioned on the bottom-left of the left eye and the upper-right of the right eye. The behavioral task was presented on a P2418HT 1920×1080p screen (Dell Inc., Texas) and was executed using MATLAB R2022b (MathWorks Inc., Massachusetts) with the Psychtoolbox-3 toolbox (Brainard, 1997).

### 2.4 Procedure

After signing the consent form, participants completed a basic demographic survey, the Anxiety Sensitivity Index 3 (Taylor et al., 2007), and the Questionnaire of Unpredictability in Childhood (Glynn et al., 2019). The last two surveys were not used for analysis in this study. Subsequently, they were fitted with an EEG cap and engaged in a classification task involving stimuli from five distinct categories presented as images or words on a computer screen.

Participants were first introduced to all the images and words from each category. All items for each category were simultaneously presented on the screen, along with the category title in the following sequence: “ANIMALS”, “TOOLS”, “FOOD”, “PLACES” (i.e., scenes), “VEHICLES”. The images and words were shown paired together on the screen so that participants had concrete examples of each word and were aware of which category each item belonged to. Notably, all words were first presented along with a concrete and specific image, suggesting that all were imaginable and concrete in the context of our task. Participants were then presented with the instructions of the task. In the task, participants were continuously presented with images and words in succession. The stimulus type alternated from trial to trial, such that images and words never followed one another. Participants were instructed to press the spacebar if the current stimulus belonged to the same category as the preceding stimulus (e.g., an image of a dog followed by the word “COW”). Otherwise (e.g., an image of a dog followed by the word “HAMMER”), no action was required. Trials requiring a response (termed “catch trials”) were balanced across category and stimulus type. These trials were designed to maintain participant engagement with the stimuli on the screen. There were no consecutive “catch trials”, meaning participants were not required to press the spacebar in back-to-back trials. Each stimulus was presented on screen for 2 s, followed by an intertrial interval of 1 *±* 0.5 s (Figure 1b).

Before beginning the task, participants completed a practice block to ensure they understood the instructions. This block consisted of 15 trials: seven images and eight words presented in alternating order, balanced across categories, with a 20% catch rate. Participants were required to repeat the practice block until they achieved a minimum score of 14/15. The main task included five blocks, each comprising 112 trials, including 10 catch trials. Blocks were planned so that 100 trials would be submitted to further analysis. The first two trials of each block were discarded, along with the catch trials. The remaining trials included all 100 stimuli, equally divided between stimulus types and categories. All catch trials, regardless of response, and all trials yielding a response, were discarded from all analyses.

Each block was completed in ∼6 minutes, and participants were provided with a score upon completion to monitor their performance. To ensure optimal data quality, participants were required to achieve a threshold score of 90% in each round for their EEG data to be included in the analysis. All participants reached this threshold in all blocks, except for a single block (score: 82.14%), which was excluded from all analyses (results were qualitatively similar when including this block). In general, results indicate that participants understood the task and were focused on it throughout the experiment. Upon completion of the procedure, the EEG cap was removed, and participants were allowed to clean up and then asked to provide feedback on the task and any additional comments they wished to share.

### 2.5 EEG preprocessing

Data was sampled at 500 Hz. EEG recordings were pre-processed using the FieldTrip Toolbox (Oostenveld et al., 2011). The data was filtered with a bandpass filter of 0.3-35 Hz and re-referenced to the average of the mastoid electrodes. We visually inspected the data and interpolated noisy channels with the data from nearby channels (1.33 ± 0.76 SD channels interpolated). All recordings underwent independent component analysis, where components containing eyeblinks, eye movements, and related artifacts were removed (2.47 ± 1.46 SD component removed). Lastly, we manually identified any artifacts in the data. The data were then segmented into trials around stimulus onset (1.25 s, 2.75 s before and after onset, respectively). Trials that included artifacts were removed from analysis (19.67 ± 19.35 SD image trials and 19 ± 19.36 word trials removed due to artifacts; no differences between word trials and image trials, *t*(29) = 1.32, *p* = 0.2). In addition, trials that included user responses were excluded. In total, out of 500 trials of data collected, we analyzed an average of 453.5 trials (± 43.8 SD) per participant (range 341–494 trials). Among the analyzed trials, there were no differences in the number of times each of the 50 words (*F*(49,1450) = 0.67, *p* = 0.96) or images (*F*(49,1450) = 0.47, *p* = 1) was presented, nor was there a difference in the number of trials across categories for words (*F*(4,145) = 0.19, *p* = 0.95) or images (*F*(4,145) = 0.15, *p* = 0.96).

### 2.6 Support vector machine classification model

We used support vector machines (SVM) to classify which category a participant was viewing using time-domain data from 58 preprocessed scalp electrodes as features (Fpz, Fp1, Fp2, AF3, AF4, AF7, AF8, Fz, F1–F8, FCz, FC1–FC6, FT7, FT8, Cz, C1–C6, T7, T8, CP1–CP6, TP7, TP8, Pz, P1–P8, POz, PO3, PO4–PO7, Oz, O1, O2), following the methods of Bae & Luck (2018). All models were implemented in MATLAB and used the *fitcecoc* function with its default settings (linear kernels, non-standardized features, no hyperparameter tuning). Each 4-s trial was divided into 182 time points, spaced 22 ms apart. Data was smoothed by averaging the data across a 62-ms window for each time point. Separate SVMs were trained at each time point, each using 58 features: the voltage measured from a single EEG channel (referenced to the mastoids). The number of features for each classifier was substantially lower than the number of trials used for most analyses to avoid overfitting.

We used a one-versus-all approach for our model, so that each SVM was trained on the 58 features to distinguish one class from the remaining classes (e.g., words vs images; images of foods vs images from all other categories). We did not tune any hyperparameters for the models and used the default settings in MATLAB. We used 5-fold cross-validation, in which an individual’s data was split into five subsets, including a training set (four subsets, 80% of the trials) and a testing set (one subset, 20% of the trials) to test the model’s accuracy. This resulted in five models, each tested on one subset and trained on the remaining four. This process was repeated 20 times, with a different set of trials divided into the five subsets in each iteration (Figure 1c). This process was repeated for each of the 30 participants at each of the 182 time points. Model accuracy was examined by comparing the predicted label to the true label at each time point. The accuracy rates for each time point were averaged across participants’ 20 iterations and five subsets.

### 2.7 Cluster-based permutation statistical analysis

We used a set of permutation tests to determine the statistical significance of our models’ performance (inspired by Bae & Luck, 2018). We generated null distributions to compare the observed models’ accuracy rates. For each experimental question (see below; e.g., classifying word trials vs. image trials), we created 1,000 randomly generated permuted datasets structured similarly to our true datasets. Permuted datasets were constructed by taking the veridical dataset and randomizing the data labels. For each participant (*N* = 30), iteration (*k* = 20), and fold (*j* = 5), the true labels were shuffled across trials. We kept labels consistent across time points for each participant to preserve autocorrelation patterns. For each permuted dataset, we calculated accuracy by comparing the model’s predictions to the shuffled labels. For both the true and shuffled datasets, accuracy rates were smoothed again across time points with an additional causal smoothing window of 9 time points (encompassing 238 ms in total), using data preceding each time point (e.g., smoothed data for time point t reflects the average of all time points between *t*-8 and *t*). We chose a causal filter to use data reflective of brain activity leading up to the time point of interest, rather than a non-causal filter which would incorporate data from future time points in the calculation. Because of this, the timing for all reported effects should be interpreted appropriately (e.g., the exact timing of the peaks depends on the specific features of the filter). Edge effects were avoided by dropping the first 8 windows, leaving 174 time windows for analyses. For both the true and shuffled datasets, we then compared the accuracy level of the model at each time point to chance levels using a two-tailed *t*-test across participants. We summed up the *t*-values for consecutive time points starting from trial onset that crossed _α_ = 0.05 (critical *t =* 2.045) to identify contiguous clusters in time that diverge from chance-level accuracy. Using the single largest cluster from each of the 1,000 permutation tests, we created a null distribution used to test the statistical significance of the true model’s accuracy levels. The summed t-values for each contiguous cluster identified in the true dataset were compared to the distribution in a one-tailed fashion, yielding its *p*-value.

For some analyses, we determined significant differences between two different models (e.g., models that used Frontal vs Parietal electrode subgroups) and used a modified version of this approach. The approach here was to test whether the difference between two true datasets was larger than that expected by chance, as reflected by a null distribution of differences between two permuted datasets. Like before, two sets of 1,000 permuted datasets were generated based on the parameters of each true dataset. For both datasets, we followed exactly the same method as above up to calculating and smoothing accuracies, using a separate permuted dataset for each of the two contrasted models. We first computed the differences between the two true datasets for each time point, yielding time-series data for this difference for each participant. The same was done for the permuted data: in each permutation, for each participant, time-series data showing the differences between the models were created. The final steps of the analysis were similar to those described above: we used two-tailed t-tests across participants to compare differences in accuracy to zero for each time point, detected contiguous clusters, and built a null distribution of differences using the permuted differences to determine the level of significance of the true differences in a one-tailed fashion.

### 2.8 Within-Subject Classification tests

The aforementioned SVM approach was used for most of our analyses. The overarching motivation for this project was to detect differences in brain activity when presented with stimuli from different categories. To detect modality-agnostic differences in categorical representations, we used all trials (both words and images) to classify across all five categories. We then investigated whether we could discern stimulus modalities by classifying between images and words. We further investigated whether EEG reflects modality-agnostic categorical representations in the brain by training SVMs on one modality (e.g., images) and testing on the other (e.g., words). In this case, we divided the data from each modality into five subsets for cross-validation, as described above, and used all five subsets for one modality’s dataset to train the model and each of the five subsets for the other modality’s dataset to test the model (e.g., image data used to train, word data used to test). This was repeated for 20 iterations of 5 blocks. Finally, we examined categorical representations for images and words separately, using image trials or word trials only.

We also examined whether different categories had more distinguishable representations by comparing categories in a pair-wise fashion (e.g., Animals vs. Food, Tools vs. Vehicles, etc.) for each modality separately. We used an alpha of 0.05 for all analyses and corrected for multiple comparisons in the pairwise analyses using the False Discovery Rate method. We next examined whether different topographical scalp regions differ in the categorical information they convey. To do this, we reran the main analyses focusing on subgroups of electrodes, examining image and word trials separately. We compared classifier accuracy between topographical regions to examine differences in representational fidelity. Since neighboring EEG electrodes tend to be strongly correlated, we repeated this analysis by sparsely selecting single representative electrodes from each of our subgroups.

Building on results from this approach, we investigated the electrodes on the left and right sides of the EEG cap separately, excluding those along the midline (odd-numbered electrodes for the left, even-numbered electrodes for the right). Classifiers were trained on each hemisphere separately for each modality. To further assess symmetry in the information reflected on each side, we trained our models on one side (e.g., left electrodes) and tested on the other (e.g., right electrodes). For this, we matched the electrode labels for each participant’s data so that the corresponding labels on each side lined up (e.g., F1 matched with F2). These models were trained over 20 iterations of 5 blocks of data shuffling, similar to our previous setup for images and words. For each block, classifiers were trained on 80% of the trials (i.e., the remaining four blocks) using data from one half of the scalp and tested on trials from the remaining block using data for the other half of the scalp.

### 2.9 Between-Subject Classification Tests

All analyses up to this point were done within-subject, with accuracy scores averaged across participants. To complement these analyses, we examined whether activity patterns were generalizable across the population (i.e., can models trained on multiple participants be successfully applied to a different participant). To test this, we adapted our analytic approach. Here, we used a leave-one-out approach, in which models were trained on data from 29 participants and tested on the remaining participant. This was repeated for all 30 participants, creating 30 datasets of time-dependent classification accuracy. Models were trained across all trials for each set of 29 participants and applied to each trial in the participant who was left out. As before, SVMs were used for this analysis. All individual trials within a given modality (e.g., words) collapsed across 29 participants were used for training, and all individual trials for the left-out participants were used for testing. Statistics were computed using the same approach described above - for each participant, labels were shuffled across all trials over 1,000 permutations, and a null distribution was created based on the permuted data. *t*-values were summed for each cluster of contiguous significant time points and compared to the null distribution, producing a *p*-value for each cluster.

### 2.10 Data and code availability

Stimuli, data, and analysis codes are available at osf.io/dfz4x.

## 3 RESULTS

### 3.1 Participant task performance

To investigate categorical representations reflected in EEG activity, 30 participants performed a task in which they viewed words and images depicting items from five categories (Animals, Food, Tools, Scenes, and Vehicles). In total, 100 visual stimuli (10 words and 10 images per category) were each presented once per block for 5 blocks, resulting in up to 500 trials for analysis. To maintain attention, participants had to respond when two stimuli belonging to the same category were presented consecutively, and they performed at ceiling (mean accuracy = 97.29%, SD = 0.19%, range 91.82% - 100%). Across all five blocks, the average score ranged from 97.06% - 97.55%.

### 3.2 Category information is represented in higher fidelity in images relative to words

We first examined whether the EEG data reflect categorical information regardless of stimulus modality (i.e., words or images). We used all trials (range 341–494 across participants) to classify between the five stimulus categories and found above-chance classification accuracy between categories (cumulative *t* = 170.43, *p* < 0.001, time range = 0–0.91 s, peak accuracy = 0.27, Figure 2a; chance level = 0.2). This demonstrated that category information is reflected in EEG topographical distributions, regardless of stimulus modality, although the effect observed in this pool dataset could still have been driven by modality-specific categorization (see Discussion). Next, we tested whether different modalities elicited distinct brain activity by training a classifier to distinguish between images and words. We indeed found above-chance classification accuracies when contrasting between modalities (cumulative *t* = 1119.8, *p* < 0.001, time range = 0–1.7 s, peak accuracy = 0.91; chance level = 0.5).

**Figure 2.**
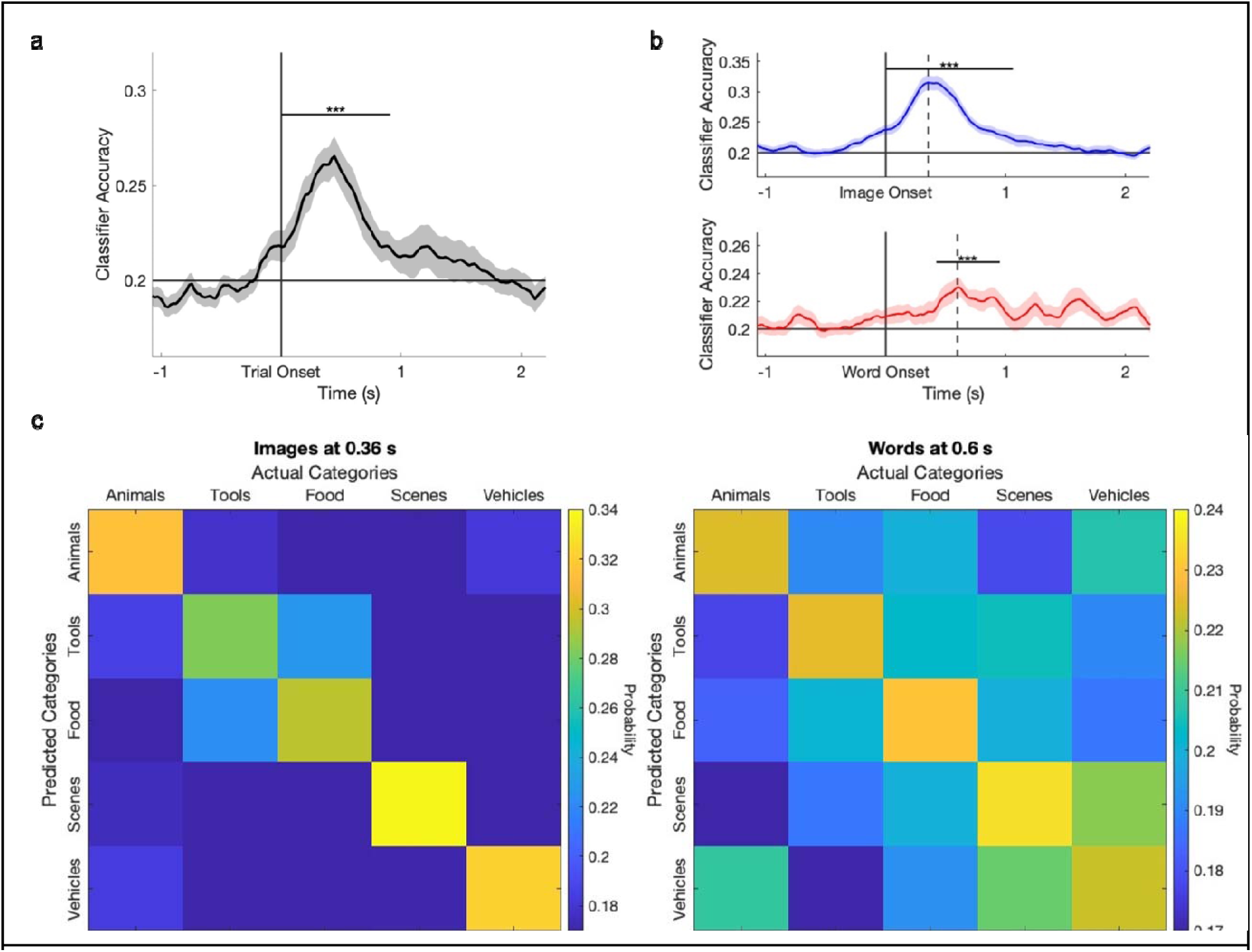
EEG neural patterns reflect category information within modality and across modalities. **a.** Classifiers trained on image- and word-trials are able to distinguish between categories. **b.** Classifiers trained on image trials (top) and word trials (bottom) can both distinguish between categories. Shaded areas reflect the standard error of the mean across participants. Dashed vertical lines mark the peak accuracy, for which the confusion matrices in panel *c* were calculated. (c) Confusion matrices, reflecting the probability of trials from each category (x-axis) to be assigned a certain category label (y-label). Probabilities were calculated for classifiers at the time at which the peak accuracy level was obtained (0.36 s for images, left panel, and 0.6 s for words, right panel) and averaged across participants. Chance level for both matrices is 0.2. **\*\*\* *p* < 0.001**

We next split our trials by modality (i.e., image and word trials separately) to examine whether modality-specific EEG data reflected category information. We found above-chance classification accuracy for images (cumulative *t* = 307.8, *p* < 0.001, time range = 0–1.06 s, peak accuracy = 0.32, trial count range = 167–247; chance level = 0.2), as well as for words (cumulative *t* = 81.64, *p* < 0.001, time range = 0.42–0.95 s, peak accuracy = 0.23. trial count range = 174–247; chance level = 0.2; Figure 2b). Confusion matrices for both images and words at their respective peak times are presented in Figure 2c. Notably, the accuracy levels obtained for images were significantly higher than for words (difference between the plots: cumulative *t* = 212.11, *p* < 0.001, time range = 0–0.82 s), showing that category information is better represented in the EEG response to visual images relative to visually presented written words. Our analytic pipeline randomly assigned trials to training and test sets, with trials including the same specific stimulus potentially used for both (e.g., of the five trials that included an image of a Cat, some may have been used for training and the other may have been used for testing). This raises the concern that classifier accuracy may be driven to some extent by item-specific rather than category-specific information. To test whether the repetition of specific items in training and testing blocks contributed to classifier accuracy, trials with a specific stimulus were randomly assigned to a specific block, which was used either for training or testing (but never both). This analytic approach, termed trial-to-block throughout the remainder of this manuscript, included random assignment was random across 20 iterations, as done for the main analysis, meaning items were randomly grouped together in blocks for each of 20 iterations (e.g., all five trials that included an image of a Cat would be included in either training or testing, but never split among the two). When controlling for trial repetition and randomizing item grouping across iterations, there were still discernible differences in EEG activity for both images (cumulative *t* = 268.45, *p* < 0.001, time range = 0 - 1.06s, peak accuracy = 0.3) and words (cumulative *t* = 79.13, *p* < 0.01, time range = 0.44 - 0.95s, peak accuracy = 0.24; chance level = 0.2), demonstrating that our classifiers are sensitive to category-specific activity rather than item-specific activity.

Finally, we examined whether modality-free category information is sufficiently reflected in the EEG data, allowing cross-modality classification. To do so, we trained classifiers on data from one modality and tested them on the data from the other modality. We found no evidence that classifiers trained on EEG responses to image presentations could be used to decode category information from word trials or vice versa. Although this may mean that the EEG activity does not reflect modality-free category information, it may also be that the classification models trained for one modality overfit to certain features that are irrelevant to the other.

We conducted additional analyses to rule out any non-category-specific influences on EEG decodability. We first examined whether the category of the stimulus in the immediately preceding trial is still discernible from the EEG activity. We did not detect any discernible activity related to the category of the previous trial (trials immediately following a word trial: cumulative *t* = 14.76, *p* = 0.32, time range = 0.33 - 0.44s, peak accuracy = 0.23; trials immediately following an image trial: cumulative *t* = 20.73, *p* = 0.18, time range = 0.73 - 0.88 s, peak accuracy = 0.23; chance level = 0.2). We also assessed the number of category transitions (e.g., Animals→Tools, Animals →Vehicles, Vehicles→Food, etc.) to consider whether some imbalance in transition probabilities may have impacted our findings, but found no difference in the transition probabilities from one category to another (*F* (19, 580) = 0.95, *p* = 0.52),

Finally, we assessed whether proximity to a catch trial or proximity to a trial yielding a user response influenced classifier accuracy. In addition to excluding catch-or response-trials, we also excluded up to two trials following a repeat/response in separate analyses for images and words. These analyses yielded results that were comparable to the main analyses, suggesting that these factors did not impact category classification (images, catch-trial proximity omission: cumulative *t* = 253.38, *p* < 0.001, time range = 0 - 0.86s, peak accuracy = 0.31; words, catch-trial proximity omission: cumulative *t* = 36, *p* < 0.05, time range = 0.42 - 0.69 s, peak accuracy = 0.23; images, response-trial proximity omission: cumulative *t* = 282.44, *p* < 0.001, time range = 0 - 1.06s, peak accuracy = 0.31; words, response-trial proximity omission: cumulative *t* = 64.2, *p* < 0.01, time range = 0.4 - 0.8s, peak accuracy = 0.24; chance level = 0.2).

### 3.3 Representation fidelity differs across categories for images

We next sought to determine which of the five categories was most distinguishable using EEG data. To test this, we trained classifiers on EEG data to distinguish between stimuli from each pair of categories (e.g., Animals vs. Tools) separately for each modality. When examining images, our results showed that all category pairs were distinguishable from one another, yielding classification accuracies above chance level (Figure 3). Categorical representations of images between Animals and Foods, Animals and Scenes, and Scenes and Tools were most distinguishable (peak accuracy rates = 0.64; chance level = 0.5), with Animals vs. Foods yielding the highest cumulative *t-*value (cumulative *t* = 234.88). These results remained significant using the trial-to-block analysis, which controls for the effects of any individual exemplar. When examining words, only one pair of categories yielded neural patterns that were significantly distinguishable (Animals and Tools; Figure 4). Notably, the timeframe for which differences were observed for this pair did not overlap with the timeframe observed when comparing across all categories (1.52 - 2.07 s vs 0.42 - 0.95 s), suggesting that this pairwise comparison might reflect a separable factor. A potential confound is that words in these categories differ in length. Indeed, words in the Animals category have a marginally lower number of average letters relative to the words in the Tools category (4.2 ± 1.72 vs. 6.3 ± 2.53; *p* = 0.06, two-sided Wilcoxon rank sum test). The late differences observed for these two categories may therefore be related to the difference in word length. When we conducted the trial-to-block analyses for these pairings, none of the pairwise comparisons were significant after correction for multiple comparisons (*p* > 0.07).

**Figure 3.**
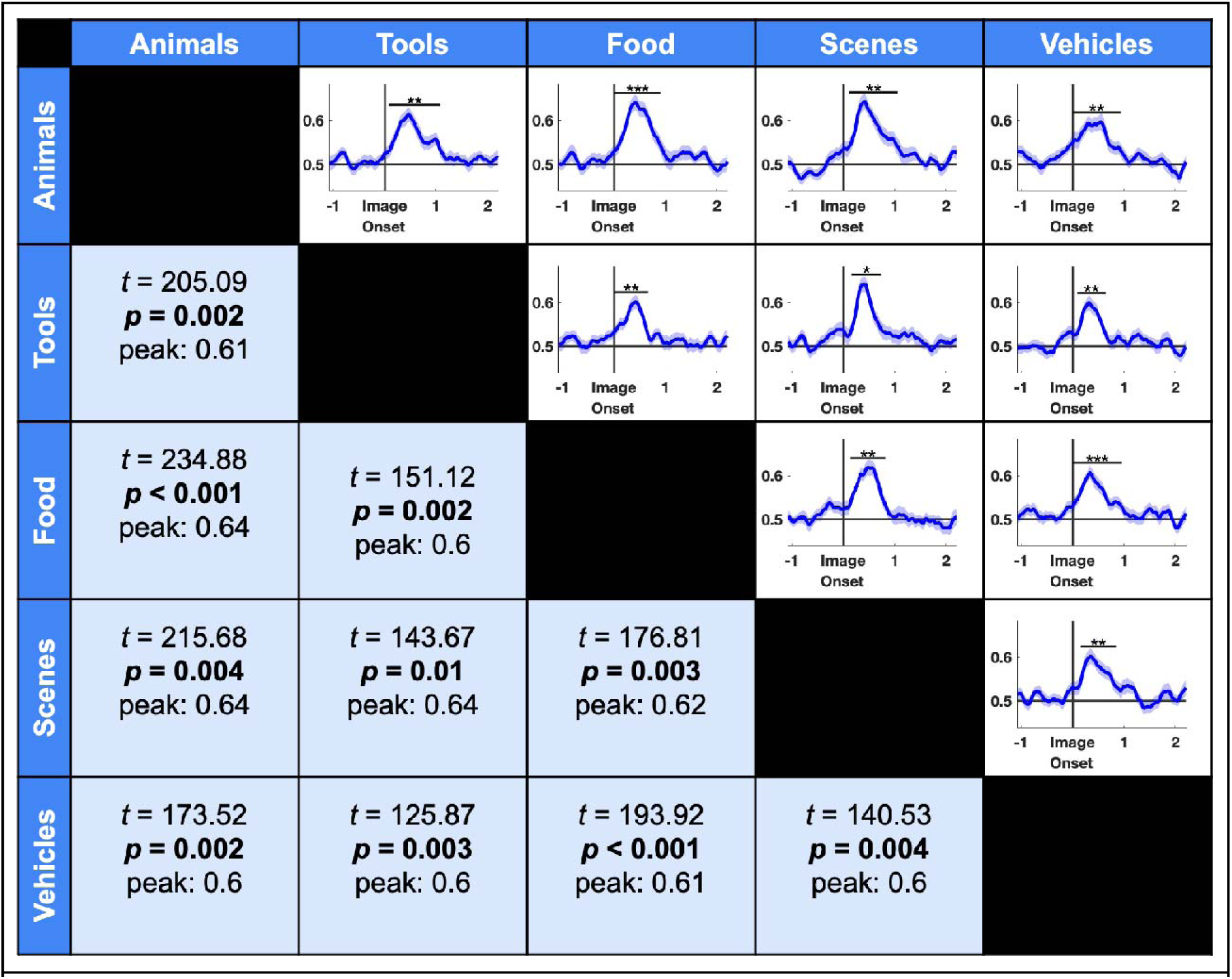
EEG data from image trials yield significant classification accuracies for all pairwise comparisons between categories. All p-values have been adjusted with the False Discovery Rate method for multiple comparisons. Shaded areas reflect the standard error of the mean across participants. Chance level accuracy is 0.5. **\* *p* < 0.05, ** *p* < 0.01, *** *p* < 0.001**

**Figure 4.**
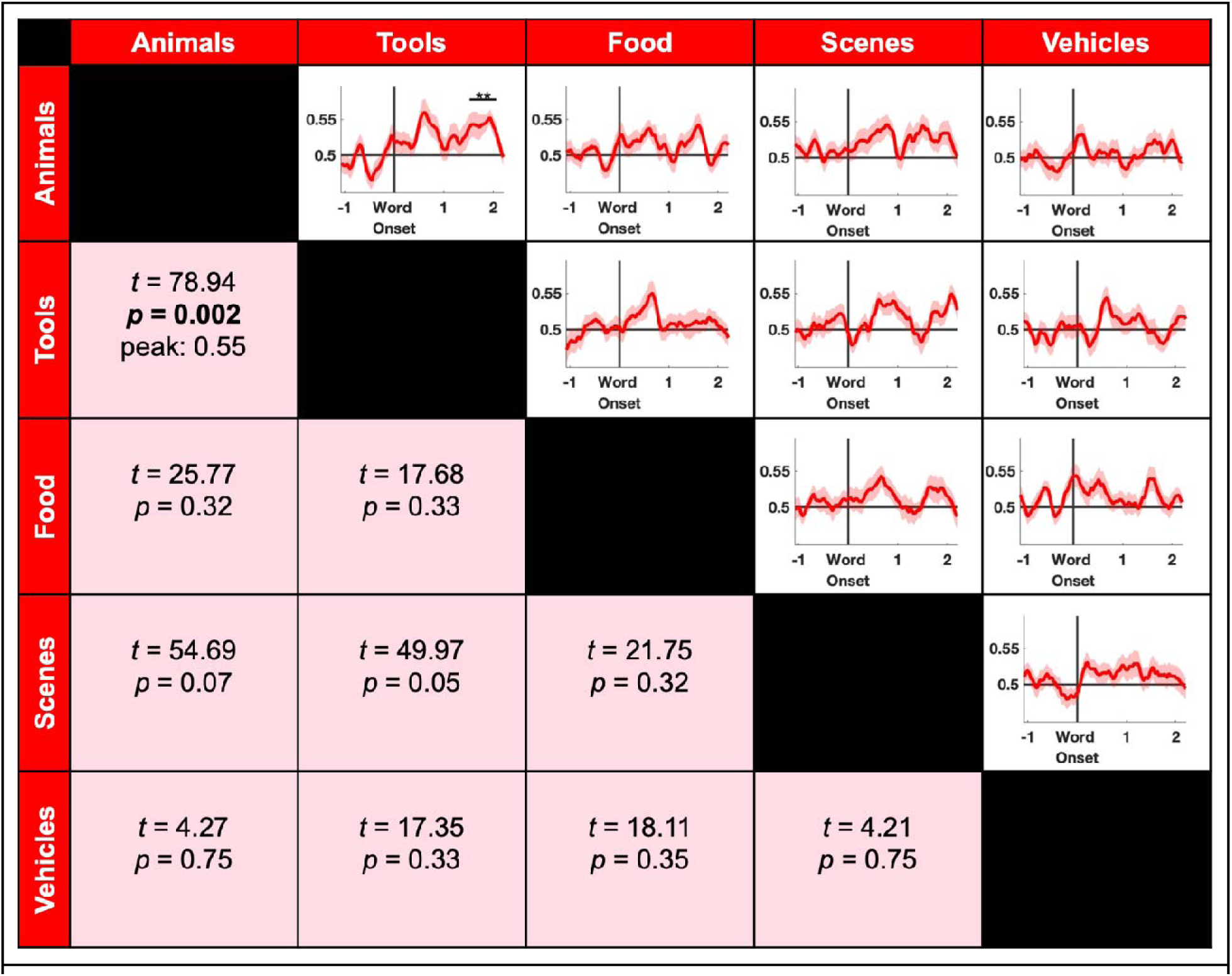
EEG data from word trials yield significant classification accuracies for one pairwise comparison between categories only (Animals vs. Tools). Non-significant *t* and *p*-values reflect the cluster closest to significance. Peak values are only shown for significant time frames. All p-values have been adjusted with the False Discovery Rate method for multiple comparisons. Shaded areas reflect the standard error of the mean across participants. Note: Animals and Tools did not remain significant in trial-to-block analyses after being corrected for multiple comparisons; see Results. Chance level accuracy is 0.5. ** *p* < 0.01

### 3.4 Information from different scalp regions differentially distinguishes between categories across modalities

We next examined how different scalp topographies contribute to cross-category classification. Although this analysis was conducted in sensor space rather than source space, we were nevertheless interested in examining the primary topographic contributors to classification accuracy across the scalp. To investigate this, we divided our electrodes into different region-specific groups: occipital, parietal, left temporal, right temporal, and frontal (Figure 5a) and trained classifiers on each subset of electrodes across all categories, split by modality. For image trials, we found that all five groups show above-chance classification accuracies: the occipital group (cumulative *t* = 267.97, *p* < 0.001, time range = 0–0.93 s, peak accuracy = 0.32; chance level = 0.2), the parietal group (cumulative *t* = 308.61, *p* < 0.001, time range = 0–1.04 s, peak accuracy = 0.32; chance level = 0.2), the left temporal group (cumulative *t* = 210.68, *p* < 0.001, time range = 0.16–1.02 s, peak accuracy = 0.28; chance level = 0.2), the right temporal group (cumulative *t* = 47.06, *p* < 0.01, time range = 0.22–0.53 s, peak accuracy = 0.21; chance level = 0.2), and the frontal group (cumulative *t* = 203.69, *p* < 0.001, time range = 0–1.04 s, peak accuracy = 0.26; chance level = 0.2). All these analyses survived FDR correction and remained significant in the trial-to-block analyses.

**Figure 5.**
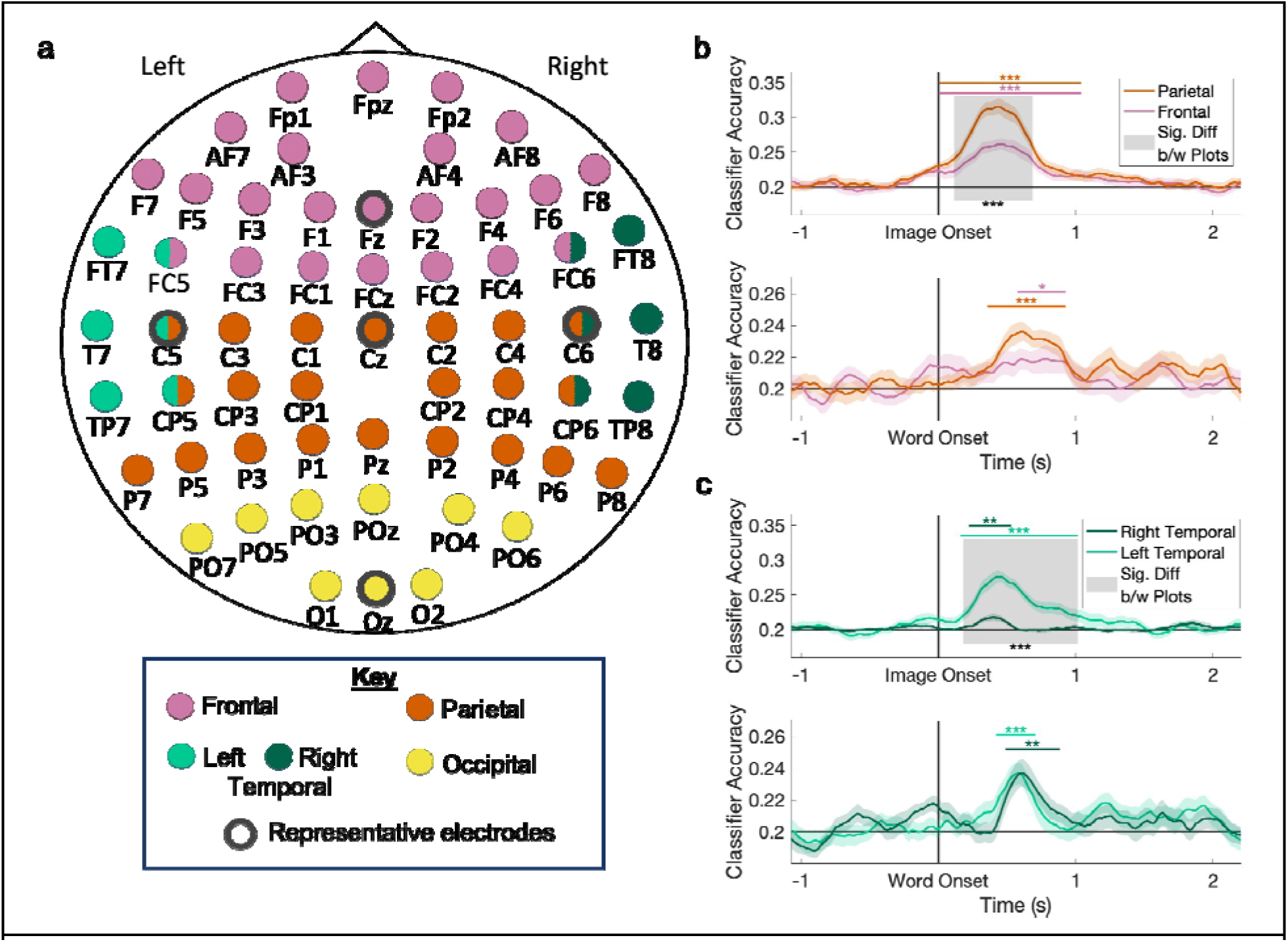
Discernible topographic contributions to classification accuracy for both words and images. **a**. The scalp electrodes were divided into groups, and each was used to train and test classifiers. Thick, grey outlines indicate which electrodes were selected to represent the subgroup in the representational electrode analyses (see text). Note that some electrodes were used in more than one group. **b.** Results showing the difference between accuracy levels obtained using the parietal electrode group (highest classifier accuracy) and the frontal electrode group (lowest classifier accuracy) for image trials (top) and word trials (bottom). Horizontal lines reflect significant time periods for each group; shaded areas represent significant differences between groups. **c.** Same as in (b), but comparing left and right temporal clusters. Shaded areas reflect the standard error of the mean across participants. Chance level accuracy is 0.2. **\* *p* < 0.05, ** *p* < 0.01, *** *p* < 0.001**

Classifier trained on word trials also yielded significant accuracies for all groups: for the occipital group (cumulative *t* = 55.18, *p* < 0.01, time range = 0.42–0.82 s, peak accuracy = 0.23; chance level = 0.2), the parietal group (cumulative *t* = 124.18, *p* < 0.001, time range = 0.36–0.93 s, peak accuracy = 0.24; chance level = 0.2), the left temporal group (cumulative *t* = 56.21, *p* < 0.001, time range = 0.42–0.71 s, peak accuracy = 0.24; chance level = 0.2), the right temporal group (cumulative *t* = 60.73, *p* < 0.01, time range = 0.49–0.88 s, peak accuracy = 0.24; chance level = 0.2), and the frontal group (cumulative *t* = 40.16, *p* < 0.05 time range = 0.58–0.93 s, peak accuracy = 0.22; chance level = 0.2). All analyses survived FDR correction. However, when using the trial-to-block analytic approach, the occipital group did not remain significant (*p* = 0.135), suggesting that classification in this region may have been driven, to some degree, by exemplar-specific patterns. To further examine the contributions of the electrode groups to classifying category information, we compared the groups yielding the highest and lowest accuracy rates (the parietal and frontal groups, respectively). For the image trials, we found that data from the parietal electrodes provided more discernible information regarding categories relative to the frontal electrodes (cumulative *t* = 128.67, *p* < 0.0011, time range = 0.11–0.69 s; Figure 5b, top). No significant differences were observed in word trials between the parietal and frontal groups (statistics for the cluster closest to significance: cumulative *t* = 21.92, *p* = 0.14, time range = 0.49–0.66 s; Figure 5b, bottom).

Data from adjacent EEG electrodes are typically correlated. Our approach for dividing the scalp into subgroups is, therefore, fundamentally limited, in that subgroups border each other and are therefore not truly separable. To overcome this limitation, we complemented this approach by examining five representative electrodes (one within each region, Figure 5a) to assess regional activity. Note that we do not claim that these distant electrodes represent independent neural generators; our motivation was primarily to identify the main scalp contributors to category classification, while remaining agnostic about the exact underlying source. Using just these five electrodes, the models are still able to classify across categories significantly above chance for both image trials (cumulative *t* =223.92, *p* < 0.001, time range = 0–0.91 s, peak accuracy = 0.28; chance level = 0.2) and word trials (cumulative *t* = 90.72, *p* < 0.001, time range = 0.42–0.95 s, peak accuracy = 0.23; chance level = 0.2). We next examined the regional contribution to category classification using these single electrodes. Despite including a single feature, this analysis still yielded classification accuracies that were significant for images across all categories (frontal, electrode Fz: cumulative *t* = 37.52, *p* < 0.01, time range = 0.0.27–0.55 s, peak accuracy = 0.21; parietal, electrode Cz: cumulative *t* = 53.16, *p* < 0.001, time range = 0.27–0.58 s, peak accuracy = 0.22; left temporal, electrode C5: cumulative *t* = 77.93, *p* < 0.001, time range = 0.27–0.69 s, peak accuracy = 0.22; right temporal, electrode C6: cumulative *t* = 35.59, *p* < 0.001, time range = 0.0.33–0.58 s, peak accuracy = 0.21; Occipital electrode, Oz: cumulative *t* = 55.67, *p* < 0.001, time range = 0.25-0.55 s, peak accuracy = 0.22; chance level = 0.2 for all). All electrodes remained significant after FDR correction. However, results for the frontal and occipital electrodes were not significant when applying the trial-to-block analytic approach. While the peaks are not far above chance, this analysis shows that image stimuli elicit EEG activity that allows for category classification with even a single electrode, though individual images may have contributed to this finding for the Fz and Oz analyses.

Word trials, however, did not produce classifiable data across all electrodes (Fz: cumulative *t* = 6.95, *p* = 0.53, time range = 0.6–0.64 s, peak accuracy = 0.21; Cz: cumulative *t* = 10.19, *p* = 0.33, time range = 1.87–1.94 s, peak accuracy = 0.21; C5: cumulative *t* = 6.65, *p* = 0.59, time range = 0.62–0.66 s, peak accuracy = 0.21; C6: cumulative *t* = 18.35, *p* = 0.14, time range = 0.53–0.66 s, peak accuracy = 0.21; Oz, cumulative *t* = 9.75, *p* = 0.36, time range = 0.58–0.64 s, peak accuracy = 0.21; chance level = 0.2 for all; all statistics for the clusters closest to significance). Interestingly, however, the right temporal electrode, C6, reached significant levels of classification in the trial-to-block analysis (cumulative *t* = 79.09, *p* < 0.001, time range = 0.44 - 0.95 s, peak accuracy = 0.24; chance level = 0.2), surviving correction for multiple comparisons. Taken together, these results demonstrate that category classification from image stimuli — and perhaps even some word stimuli — is feasible even when using low-density EEG systems.

We next considered the left and right temporal subsets of electrodes separately to assess whether they asymmetrically captured category information. Interestingly, accuracy levels were significantly higher for the left cluster relative to the right one (cumulative *t* = 185.18, *p* < 0.001, time range = 0.18–1.02 s; Figure 5c, top). There were no differences between the two subgroups for words (Figure 5c, bottom). Overall, results revealed varying levels of discernible EEG activity across different scalp areas. These differences were more pronounced for image stimuli, which also elicited asymmetrical activity between the left and right temporal regions.

We next examined the lateralized contributions to category classification more directly by dividing all electrodes into left- and right-scalp subgroups, omitting midline electrodes. We first tested whether each half was sufficient to classify between categories, and found above-chance accuracy levels for both left and right scalp electrodes for image trials (left scalp: cumulative *t* = 295.01, *p* < 0.001, time range = 0– 1.19 s, peak accuracy = 0.27; right scalp: cumulative *t* = 259.44, *p* < 0.001, time range = 0–0.99 s, peak accuracy = 0.28; chance level = 0.2). For word trials, only data from the right scalp electrodes yielded accurate classifiers (right electrodes: cumulative *t* = 69.05, *p* < 0.001, time range = 0.44–0.93 s, peak accuracy = 0.22; left electrodes: cumulative *t* = 27.22, *p* = 0.05, time range = 0.49–0.66 s, peak accuracy = 0.22; chance level = 0.2). Left and right scalp activity were not significantly different from each other for either modality (images, statistics for the cluster closest to significance: cumulative *t* = 21.19, *p* = 0.26, time range = 0.18–0.33 s; words, statistics for the cluster closest to significance: cumulative *t* = 10.19, *p* = 0.4, time range = 1.52–1.59 s; chance level = 0.2).

We further investigated the neural symmetry of information representation by training classifiers on one group of electrodes (e.g., those over the left hemisphere) and testing on the other (e.g., those over the right hemisphere). The objective of this analysis was to determine whether category-specific information was symmetrically represented across the two sides of the scalp for images and words. Indeed, we found evidence for symmetry. For both image and word trials, we found above-chance classifier accuracy when the right electrodes were used for training and the left for testing (images: cumulative *t* = 40.12, *p* < 0.05, time range = 0.33–0.62 s, peak accuracy = 0.21; words: cumulative *t* = 32.83, *p* < 0.05, time range = 0.49–0.71 s, peak accuracy = 0.21; chance level = 0.2). When left electrodes were used for training, testing on the right electrodes for image trials produced marginally significant accuracy rates (cluster closest to significance: cumulative *t* = 26.71, *p* = 0.06, time range = 0.36-0.55 s, peak accuracy = 0.21; chance level = 0.2). For the word trials, however, training on the left electrodes and testing on the right ones did not yield above-chance accuracy rates (cluster closest to significance: cumulative *t* = 4.25, *p* = 0.72, time range = 1.5-1.52 s, peak accuracy = 0.21; chance level = 0.2). Significance levels did not change after FDR correction, except for the marginal effect for the classifiers trained on the left hemisphere for images, which was no longer marginally significant.

### 3.4 Between-subjects classification

We next tested whether the category-specific EEG topographies were generalizable across participants. Up to this point, classifiers were trained for each participant independently. Conceivably, each participant’s time-dependent and category-specific neural patterns may be entirely idiosyncratic. However, there may be commonalities across participants. To test this, we examined whether classification models can be successfully applied to participants other than the one on which they were trained. Using a leave-one-out approach, we found that when the models were trained and tested on images, we were able to generalize time-dependent models across participants, although accuracy levels were notably lower than those produced by within-participant models (cumulative *t* = 62.03, *p* < 0.05, time range = 0–0.38 s, peak accuracy = 0.211; chance level = 0.2; Figure 6, top). However, models trained and tested on word trials did not generalize across participants (statistics for the cluster closest to significance: cumulative *t* = 13.74, *p* = 0.33, time range = 1.02–1.1 s, peak accuracy = 0.21; chance level = 0.2; Figure 6, bottom). These results suggest that category-specific neural patterns reflected in EEG data are generalizable across the population for image representations of categories, but the magnitude of this effect is small, going only slightly above chance levels.

**Figure 6.**
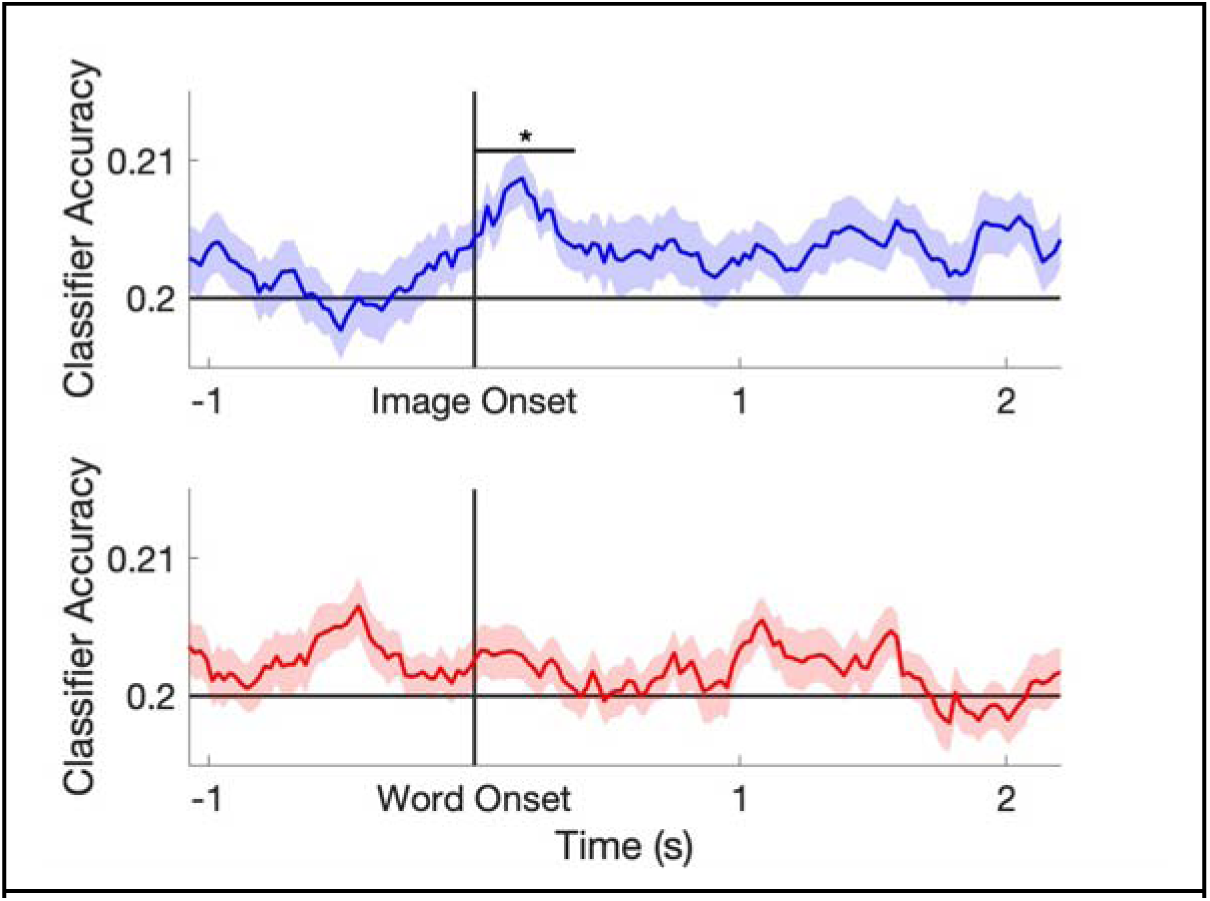
Results from the cross-participant classification analysis show above-chance accuracy for images (top) but not for words (bottom). Shaded areas reflect the standard error of the mean across participants. Chance level accuracy is 0.2. * *p* < 0.05

## 4 DISCUSSION

The goal of this study was to build upon recent methodological advances to demonstrate the potential of EEG for studying category-level neural representations across modalities. Using high-density EEG, we found discernible category representations, as well as image-and word-specific activity patterns. When comparing across modalities, EEG data following image presentation reflected more category information than data collected following word presentations, yielding higher classification accuracy. Among the five examined categories, pairwise contrasts differed in the fidelity of the trained classifiers, suggesting that some categories are more discernible from the scalp data. We further examined whether specific topographical regions contributed differentially to classifier accuracy, and found the highest accuracy rates for electrodes positioned in parietal regions. There were also significant differences in classifier accuracy between left and right temporal electrodes, with left electrodes reflecting more category information relative to right temporal ones. Similar results were obtained using single representative electrodes, demonstrating that category classification is achievable with low-density EEG as well. Lastly, we show that category-specific EEG topography is generalizable across the population using a leave-one-out approach across participants. Taken together, these results demonstrate that EEG can be used to investigate the neural representations of categorical information.

Although our main focus in this study was methodological, our results provide insight into how the brain represents categories in modality-specific and modality-agnostic ways. A key feature of our experimental design is that we contrasted data collected in trials where images were presented and data collected in trials where words were presented. Analyzing each modality separately demonstrated that data from image trials yielded higher classification accuracies than data from word trials. More broadly, across multiple classification schemes and research questions, image trials far outperformed word trials. Notably, word trials still yielded above-chance classification accuracy, demonstrating that our models are able tap into true category representations rather than perceptual differences. Collapsing over modalities, we were able to classify across categories. However, classification accuracy for the pooled dataset may be driven by the subgroup of training and testing trials belonging to one modality rather than a modality-agnostic representation. Indeed, we were unable to successfully train a classifier on one modality and apply it to the other. In this case, as in all other analyses utilizing the word trials, it is impossible to tell whether these null results are due to inherent limitations of EEG in reflecting higher-order cross-modal representations, or due to power issues borne out of the relatively low accuracy rates for word trials (peak = 0.23; chance level = 0.2).

Early attempts to investigate category representations using neuroimaging tools utilized positron emission tomography (Martin et al., 1996), and later fMRI, primarily due to their superior spatial resolution and access to neural sources throughout the brain. However, fMRI is limited due to its relatively high costs, which frequently lead to underpowered designs (Turner et al., 2018). In contrast, EEG is a more affordable neuroimaging method (especially for low-density systems). Other advantages of EEG over fMRI are that it is more tolerable to movement, facilitating more naturalistic designs and recording during sleep. Finally, EEG has better temporal resolution, which provides information about the rapid temporal dynamics of the activity of category neural representations. Traditionally, EEG research on category representations focused on ERPs (e.g., N170 for faces; Bentin et al., 1996). With recent methodological progress, EEG-based machine learning approaches have become increasingly popular. In this manuscript, we leverage these recently employed techniques and the unique features of EEG to demonstrate its ability to identify category-specific neural activation patterns, which could lend itself to future research on category-selective neural activity. Although machine learning has been used to investigate categorical representations in the past (e.g., Schreiner et al., 2021), our work is unique in its comprehensive examination of multiple categories across modalities.

To determine which category pairs were more distinctly represented in the EEG signal, we conducted pairwise comparisons for each modality and found that all category pairs for image trials and one pair for word trials (Animals vs. Tools) elicited significantly distinguishable activity. Whereas the differences for images were robust, the pairwise differences for words were scarce and less reliable. Notably, the time span for the significant difference between words in the Animals vs Tools classifier was later than the time span for all categories of words (1.52 - 2.07 s vs.0.42 - 0.95 s), and this pairing did not remain significant in the trial-to-block analysis, suggesting that this effect may be separable from the cross-category differences identified in the analyses incorporating all five categories. None of the word-trial pairwise comparisons showed a significant difference in the earlier time frame, where differences were observed when using all five categories, and this likely stems from the small effect size (peak accuracy for words = 0.23, chance level = 0.2). Among the category pairs for images, the classifier distinguishing the Animal and Food categories performed best. Future experimental designs intended to leverage EEG for cross-category classification are therefore encouraged to use these two categories.

To examine topographical contributions to the EEG-detection of brain activity, we separated our electrodes into subgroups for classifier training. This analysis was not intended to identify the neural sources of different categories, but to estimate the contribution of different topographical locations on the scalp to guide future EEG studies. We found that all our subgroupings reflected category information for both image and word trials. When using the trial-to-block analytic approach, the occipital region for words was no longer classifiable, suggesting electrodes in that subregion relied more on data from specific item repetition than categories in the original analyses. Specifically, the parietal subgroup performed best in both modalities, whereas the frontal subgroup performed worst. To examine isolated electrode activity and demonstrate the feasibility of using low-density EEG setups, we reduced these subgroups to single representative electrodes and found that classification accuracy remained high when examining just five electrodes (along with the mastoid electrodes, which acted as references). Data from individual electrodes was sufficient for classifying categories in image trials but not in word trials.

Finally, we extended our results by considering across-participant category representations. All analyses mentioned up until this point were agnostic as to whether the neural representations of categories were generalizable across the population; they only considered the reliability of within-participant patterns across stimuli. For this analysis, we directly examined the generalizability of the neural representations across participants. Since different classifiers were trained at each time point, above-chance across-participant classification accuracy would require both generalized category-specific representations and common temporal dynamics. Our results demonstrate above-chance classification using data collected in image trials but not word trials. As before, it is unclear whether this null effect is due to some inherent property of words or a result of the upper bound on classification accuracy for this modality, limiting statistical power. Regardless, the findings showing above-chance classification for images demonstrate that there are indeed some generalizable representations, although the effect size was small (peak accuracy = 0.211, chance level = 0.2). The small effect demonstrates the limited potential of the cross-participant approach and highlights its limitations relative to the better-performing within-participant classifiers trained on idiosyncratic category-specific neural patterns.

Our study, and indeed many studies utilizing EEG, have some notable limitations. The biggest limitation is that the neural origin of the category representations is unclear. The neural underpinnings reflected in EEG signals are notoriously hard to localize, and our results are impacted by this limitation. Although data collected from the scalp indicate that there are category-specific differences in neural activity (and reveal their temporal dynamics), they leave open the question of which brain networks drive the observed differences. Another limitation is specific to our study; in some analyses, accuracy levels perplexingly seem to increase before image onset. We have confirmed that it is not due to trigger delays in our system and therefore believe that this may be due to the zero-phase lowpass filter used in our preprocessing code, which smooths the data in a non-causal way. To address this issue, we only considered data after image onset for all analyses. A final limitation concerns potential confounds that may partially explain the differences captured by the neural decoders. Our choice of word stimuli was intended to minimize alternative explanations, such as differences in word frequency and length between categories. Furthermore, we attempted to render all words concrete and imaginable by presenting them along with their corresponding images. Nevertheless, there may still be some differences between categories beyond each word’s category affiliation. Similarly, there may be some low-level visual differences between images in different categories that may have contributed to some degree to classifier performance.

In this manuscript, we have presented an experimental design and analytic pipeline for identifying category representations using EEG data. Our data and approach could easily be adopted to address multiple research questions. Importantly, detecting category representation using EEG would allow for monitoring of neural reactivation during offline states (e.g., sleep or wakeful rest), whether accompanied by conscious perception or not (Tal & Schechtman, 2025). Our approach, therefore, supplements the current toolkit used in cognitive neuroscience to examine the manner in which covert or overt neural representations support cognitive operations.

## Conflict of interest statement

The authors declare no competing financial interests.

## Acknowledgments

This work is supported by the National Institute of Health (USA) grant R00—MH122663. The authors wish to thank members of the Cognitive Neuroscience of Sleep lab, and especially Matthew Cho, for their assistance with data collection and feedback on study design.

## Notes

https://osf.io/dfz4x

### Competing Interest Statement

The authors have declared no competing interest.

### Summary of Updates

Revision after reviewer comments, including mostly results section

